# Carry on caring: infected females maintain their level of parental care despite suffering high mortality

**DOI:** 10.1101/2020.09.10.291401

**Authors:** Tom Ratz, Katy M. Monteith, Pedro F. Vale, Per T. Smiseth

## Abstract

Parental care is a key component of an organism’s reproductive strategy that is thought to trade-off with allocation towards immunity. Yet it is unclear how caring parents respond to pathogens: do infected parents reduce their amount of care as a sickness behaviour or simply from being ill, or do they prioritise their offspring by maintaining high levels of care? Here we explored the consequences of infection by the pathogen *Serratia marcescens* on mortality, time spent providing care, reproductive output, and expression of immune genes of female parents in the burying beetle *Nicrophorus vespilloides*. We compared untreated control females with infected females that were inoculated with live bacteria, immune-challenged females that were inoculated with heat-killed bacteria, and injured females that were injected with buffer. We found that infected and immune-challenged females mounted an immune response and that infected females suffered increased mortality. Nevertheless, infected and immune-challenged females maintained their normal level of care and reproductive output. There was thus no evidence that infection led to either a decrease or an increase in parental care or reproductive output. Our results show that parental care, which is generally highly flexible, can remain remarkably robust and consistent despite the elevated mortality caused by infection by pathogens. Overall, these findings suggest that infected females maintain a high level of parental care; a strategy that may ensure that offspring receive the necessary amount of care but that might be detrimental to the parents’ own survival or that may even facilitate disease transmission to offspring.

## Introduction

When infected by a pathogen, animals often alter their behaviours and social interactions (Hart, 1988; Kelley et al., 2003; Adelman & Martin 2009; Vale et al., 2018). This change in behaviour may occur as a side effect of lethargy (Adelman & Martin, 2009) or it may represent what is known as sickness behaviour; a strategic decision to shift resources towards immune defence by reducing activity levels (Lopes et al., 2016; van Kerckhove et al., 2013) and costly social interactions (Bos et al., 2012). Lethargy may be a consequence of the pathogen negatively impacting on the host’s ability to remain active, thus leading to reduced mobility (e.g. Bradley et al., 2005; Cameron et al., 1993), foraging (e.g. Levri & Lively, 1996; Venesky et al., 2009) and social activity (Lopes et al., 2016). Lethargy may also be associated with sickness behaviour, an adaptive adjustment to fight the infection that allows the host to diverge resources from non-essential activities, such as social interactions, to the immune system (Hart, 1988; Exton, 1997; Johnson, 2002). When individuals interact with family members, sickness behaviour may also help reduce the risk of disease transmission to close kin (Heinze & Walter, 2010; Stroeymeyt et al., 2018) as a possible kin-selected behaviour (Shakhar & Shakhar, 2015; Shakhar, 2019). However, recent empirical evidence shows that sick individuals often maintain their social interactions with close kin (Lopes et al., 2018; Stockmaier et al., 2020). Yet empirical studies testing the effects of infection on social behaviour towards close kin are still scarce, with most studies being based on immune challenges (injecting with heat-killed pathogens or products from pathogens; e.g. Aubert et al., 1997; Bonneaud et al., 2003; Stockmaier et al., 2020) that exclude potential effects of the pathogen on host’s behaviour.

Parental care is a key component of an organism’s reproductive strategy in many birds, mammals, and insects (Royle et al., 2012) that is thought to trade-off with allocation of resources towards immunity (Richner et al., 1995). Caring parents incur costs of care in terms of increased energy expenditure, reduced opportunities for additional reproductive attempts, reduced survival, and/or reduced future reproductive success (Williams, 1966). Parental care enhances offspring growth and/or survival by neutralising environmental hazards to offspring, including risks associated with starvation, predation, parasitism, and competition (Royle et al., 2012). Thus, when infected by a pathogen, parents face the dilemma of whether to shift allocation towards immunity at the expense of maintaining their level of parental care, or maintain the level of parental care at the expense of increasing their allocation towards immunity. Parents that reduce their level of care to increase their immune response would risk impairing their offspring’s growth and survival, whereas parents that maintain their level of care would risk falling ill by not mounting an adequate immune response. Experimental studies using immune-challenges found that female laboratory mice tend to maintain their level of care and maintain normal offspring growth and survival (Aubert et al., 1997), while house sparrows drastically reduce their food provisioning at the cost of reduced offspring survival (Bonneaud et al., 2003). Thus, it is unclear how caring parents balance allocation towards parental care and immunity in response to infection: do infected parents reduce or maintain their level of care, and is there a trade-off between the magnitude of the immune responses and the level of parental care?

Here, we investigated how parents balance their allocation towards parental care and immunity in response to infection in the burying beetle *Nicrophorus vespilloides.* This is an ideal system to investigate this issue because it is one of the few insects with extensive parental care. Parental care includes provisioning of food to larvae, defence against predators and infanticidal conspecific intruders and production of antimicrobials and enhances the offspring’s growth and survival (Scott, 1998; Eggert et al., 1998; Smiseth et al., 2003; Rozen et al., 2008). Burying beetles show changes in immunity during parental care (Steiger et al., 2011), which include differential expression of antimicrobial peptides (Jacobs et al., 2016; Ziadie et al., 2019). Parents may mount a personal immune response that helps them deal with pathogens. However, there is also evidence that parents invest in social immunity that benefits the offspring but is costly to the parents (Cotter & Kilner, 2010b; Ziadie et al., 2019). Social immunity in burying beetles occurs as parents coat the carcass with exudates with potent antibacterial activity (Cotter & Kilner, 2010b), which reduces microbial load and improves the offspring’s survival (Rozen et al., 2008).

To test for a causal effect of infection on parental care and immunity, we monitored the amount of care provided by infected females that were inoculated with live bacteria, immune-challenged females that were inoculated with heat-killed bacteria, injured females that were injected with buffer, and untreated control females. We also monitored their life span and overall reproductive output. In parallel, we quantified the personal and social immune responses of females in each treatment by measuring the expression of genes encoding antimicrobial peptides, namely *attacin-4, cecropin-1, coleoptericin-1* and *PGRP-SC2.* If females respond to infection by shifting their allocation towards immunity, we would expect infected and/or immune-challenged females to show a reduction in parental care and an increase in the overall expression of immune genes. Alternatively, if females respond to infection and/or immune-challenges by shifting allocation towards current reproduction, we would infected and/or immune-challenged females to maintain their level of parental care and show a reduction the overall expression of immune genes. Assuming there is a trade-off between personal and social immunity (Cotter & Kilner, 2010a), we expect an increase in the expression of genes involved in personal immunity relative to the expression of genes involved in social immunity if infected and/or immune-challenged females shift allocation towards their own immunity. Alternatively, we would expect a reduction in the expression of genes involved in personal immunity relative to the expression of genes involved in social immunity if infected and/or immune-challenged females shift allocation towards current reproduction.

## Materials and methods

### Origin and rearing of experimental beetles

Experimental beetles originated from wild individuals collected in the Hermitage of Braid and Blackford Hill Local Nature Reserve, Edinburgh, U.K. The beetles had been maintained in a large outbred population (200–300 individuals were bred per generation) under laboratory conditions for at least 5 generations before the start of our experiment. Non-breeding adult beetles were housed in individual transparent plastic containers (12 cm x 8 cm x 2 cm) filled with moist soil, under constant temperature at 20°C, 16:8h light:dark photoperiod and ad libitum access to organic beef as food supply.

### Experimental design and procedures

To investigate the effects of infection on parental care, reproductive output and immunity, we used a group of untreated control females (*N*_Control_ = 61) and three groups of experimental females: infected females that were inoculated with the pathogenic bacteria *Serratia marcescens* (*N*_Infected_ = 58), immune-challenged females that were inoculated with heat-killed bacteria (*N*_Challenged_ = 70), and injured females that were injected with buffer (*N*_Injured_ = 56). At the beginning of the experiment, each individual virgin female was randomly assigned an unrelated male partner and transferred to a larger plastic container (17 cm x 12 cm x 6 cm) lined with moist soil and containing a freshly thawed mouse carcass of a standardized size (19.97–23.68g) (Livefoods Direct, Sheffield). We weighed each female on the day before the anticipated hatching date (i.e. two days after the onset of egg-laying; Smiseth, Ward, & Moore, 2006). We then placed females in an individual plastic vial plugged with cotton. Females remained in this vial until we applied the treatment (see details below), after which they were transferred into a new large container containing fresh soil and supplied with their original carcass. We left the eggs to develop in the old container, while males were discarded. We separated the females from the eggs so that we could allocate each female with an experimental brood of 15 same-aged larvae of mixed maternal origin. We removed the male to avoid any potential effects of male parental care buffering against effects of the experimental treatment on the female. Male removal has no effect on the developing brood under laboratory conditions (Smiseth et al., 2005). We next set up experimental broods of 15 larvae by collecting newly hatched larvae emerging in the soil, starting the day following the separation of females and eggs. We generated experimental broods by pooling larvae that had hatched from eggs laid by multiple females (Smiseth et al., 2007). We used a standardized brood size that was comprised of 15 larvae of a known age to avoid any potential confounding effects of variation in the number and age of the larvae on maternal behaviour (Smiseth et al., 2003; Ratz & Smiseth, 2018). Given that parents will kill any larvae that emerge on the carcass before their own eggs have hatched (Müller & Eggert, 1990), we only allocated an experimental brood to a female once her own eggs had hatched.

### Bacterial preparation

We chose *Serratia marcescens* (strain DB11) as an appropriate natural bacterial pathogen for *N.vespillodies. Serratia marcescens* is a gram-negative bacterium commonly found in the soil and on decomposing carrion (Hejazi & Falkiner, 1997; El Sanousi et al, 1987). It has been shown to infect several insect species and is known to cause mortality in both eggs and larva of *N. vespilloides* (Wang & Rozen, 2018; Jacobs et al, 2014). Pilot tests confirmed that *S. marcescens* increased female mortality (Ratz et al., unpublished data), but only when injected above a certain concentration and volume (see below). We also note that our pilot tests showed that stabbing with *Pectobacterium carotovorum, Pseudomonas aeruginosa,* and injections with *Pseudomonas entomophila* had no detectable effect on female mortality.

To grow the *S. marcescens* culture, we inoculated 10 mL of Luria-Bertani (LB) broth (Fisher Scientific) with 200 μL of a frozen 25% glycerol suspension from a single isolated *S. marcescens* colony. The culture was aerobically incubated overnight in an orbital shaker at 140 rpm and 30°C. On the day of infection, the overnight culture was diluted 1:10 into fresh LB broth and incubated under the same conditions until the culture had reached the mid-log growth phase (OD_600_ 0.6–0.8). Optical density was checked using a microplate absorbance reader at an absorbance of 600 nm. The mid-log phase culture was pelleted by centrifugation (15 min, 4°C, 2500 rpm) and the supernatant removed. The pellet was then re-suspended in sterile Phosphate Buffer Saline (PBS, pH 7.4) and adjusted to OD_600_ 1. The final inoculum OD_600_ was calculated as described in Siva-Jothy et al. (2018). The final inoculum was split into two tubes; one tube was heated to 70°C for 45 min killing the bacteria and allowing for an immune-challenged treatment group while the other tube was kept as a live culture for the infected treatment group.

### Infection procedure

On the day preceding the expected date of hatching, we randomly allocated each female to an experimental treatment group. Female from all treatment groups were first anesthetised by releasing CO2 into their individual tube for 40 s. Control females were then returned to their vials to recover for 30 min, while experimental females were placed on a CO2 pad under a dissecting microscope. We used a glass needle attached to a microinjector (Nanoject II, Drummond Scientific Co) to inject injured females with 0.552 μL of sterile PBS buffer, immune-challenged females with 0.552 μL of heat-killed *S. marcescens* solution, and infected females with 0.552 μL of OD_600_ 1 live *S. marcescens* solution (~1.3 million colony forming units). We performed the injection by introducing the needle through the soft cuticle that joins the thorax and the abdomen on the ventral side (Reavey et al., 2014). Once injected, experimental females were returned to their vials to recover for 30 min. Following recovery, we next moved control and injected females back to the large containers containing their carcasses.

### Maternal care, female weight change, female mortality, and offspring performance

We recorded the amount of care provided by each female 24 h (±15 min) after we placed the larvae on the carcass, which corresponded to 48 h (±4 h) after females were handled and/or injected. We performed direct observations under red light for 30 min, recording maternal behaviour every 1 min in accordance with established protocols (e.g., Smiseth & Moore 2002, 2004; Ratz & Smiseth 2018). We recorded maternal care as food provisioning, defined as when there was mouth-to-mouth contact between the female and at least one larva, and carcass maintenance, defined as when the female was excavating the soil around the carcass or coating the carcass with antimicrobial secretions. We conducted the behavioural observations blindly with respect to treatment, as it was not possible for the observer to identify the experimental treatments.

Females and their broods were then left undisturbed until larvae completed their development, at which stage they left the mouse carcass to disperse into the soil. At dispersal, we weighed the female, counted the number of larvae and weighed the brood. We estimated weight gain over the reproductive attempts by the female as the difference in body mass between egg-laying and larval dispersal. We estimated larval survival as the difference between the final brood size at dispersal and the initial brood size at hatching (i.e. 15 larvae), and mean larval mass as the total brood mass divided by brood size.

### Hemolymph sampling, RNA extraction, reverse transcription, and qPCR

To examine the effects of the treatment on the female’s immune response, we quantified the expression of genes coding for antimicrobial peptides (AMPs) by quantitative real-time polymerase chain reaction (qRT-PCR). We focused on the expression of the four following genes: *attacin-4, cecropin-1, coleoptericin-1* and *PGRP-SC2.* We focused on these genes because they are known to have a role in the immune response against gram-negative bacteria, such as *S. marcescens* (Imler & Bulet, 2005; Vilcinskas et al., 2013a,b) and there is some knowledge about their function in personal or social immunity in this species (Jacobs et al., 2016; Parker et al., 2015; Ziadie et al., 2019): *attacin-4, cecropin-1,* and *coleoptericin-1* seem to play a role mainly in personal immunity (Jacobs et al., 2016), while *PGRP-SC2* plays a role in social immunity (Parker et al., 2015; Ziadie et al., 2019).

In parallel with the behavioural observation, we randomly selected a subset of females for RNA extraction, which included 13 control, 14 injured, 17 immune-challenged, and 14 infected females. We removed each of these females from their containers 48 h (±4 h) after infection, and placed them in an individual plastic vial plugged with cotton. We then anesthetised each female with CO2 as described above. Once anesthetised, we extracted hemolymph from each female placed on a CO2 pad by puncturing the soft cuticle behind the thorax with a micro-pine and then drawing hemolymph with a 10 μL-glass capillary. We sampled 2 μL to 10 μL of hemolymph per female and transferred it into 1.5μl-micro-tubes containing 100 μL of TRIzol reagent (Invitrogen, Life Technologies). All hemolymph samples were then stored at −70°C until RNA extraction.

RNA extractions were performed using the standard phenol-chloroform method and included a DNase treatment (Ambion, Life Technologies). The RNA purity of eluted samples was confirmed using a Nanodrop 1000 Spectrophotometer (version 3.8.1). cDNA was synthesized from 2 μL of the eluted RNA using M-MLV reverse transcriptase (Promega) and random hexamer primers, and then diluted 1:1 in nuclease free water. We performed quantitative RT-PCR on an Applied Biosystems StepOnePlus machine using Fast SYBR Green Master Mix (Applied Biosystems). We used a 10 μL reaction containing 1.5 μL of 1:1 diluted cDNA, 5 μL of Fast SYBR Green Master Mix and and 3.5 μL of a primer stock containing both forward and reverse primers at 1 μM suspended in nuclease free water (final reaction concentration of each primer 0.35 μM). For each cDNA sample, two technical replicates were performed for each set of primers and the average threshold cycle (Ct) was used for analysis.

Primers were designed based on amino acid sequences provided on Kyoto Encyclopedia of Genes and Genomics (KEGG) or supplementary information provided by Jacobs et al. (2016) (KEGG: PGRP-SC2, Rlp7; Jacobs et al. 2016: Attacin-4, Coleoptericin-1, Cecropin-1). Briefly, the amino acid sequence was entered into the Basic Local Alignment Search Tool (BLAST) on NCBI.gov, the accession number producing the most similar alignments within *N. vespilloidies* was selected and the corresponding nucleotide sequence used for primer design in Primer3 (version 4.1.0) and Beacon Designer (Premier Biosoft International). All primers were obtained from Sigma-Aldrich Ltd; Attacin-4_Forward: 5’ GCATTTACACGCACAGACCT 3’, Attacin-4_Reverse 5’ CGGCAACTTTACTTCCTCCG 3’; Cecropin-1_Forward 5’ CGAGCACACAACAGTTCCTT 3’, Cecropin-1_Reverse 5’ ATCAAAGCTGCGATGACCAC 3’; Coleoptericin-1_Forward 5’ GAAACGGTGGTGAACAGGTG 3’, Coleoptericin-1_Reverse 5’ GAGTCTTGGGGAACGGGAA 3’; PGRP-SC2_Forward 5’ CGAAGGTCAAGGTTGGGGTA 3’, PGRP-SC2_Reverse 5’ GTTCCGATGACACAGATGCC 3’. We used Rpl7 as an endogenous reference gene, following Jacobs et al. (2014, 2016) and Cunningham et al. (2014); Rpl7_Forward 5’ GTCGGCAAGAACTTCAAGCA 3’, Rpl7_Reverse 5’ TCCCTGTTACCGAAGTCACC 3’. For each pair of primers the annealing temperature (TO) was optimised and the efficiency (Eff) of each primer pair calculated by 10-fold serial dilution of a target template (each dilution was assayed in duplicate); Attacin-4: T□= 59°C Eff= 102.21%, Cecropin-1: T□= 59.5°C Eff= 102.26%, Coleoptericin-1 T□= 61.6°C Eff= 101.86%, PGRP-SC2: T□= 60.2°C Eff= 99.84%, Rpl7: T□= 60°C Eff= 98.25%.

### Statistical analysis

All statistical analyses were conducted using R version 3.6.0 (R Development Core Team, 2019) loaded with the packages *car* (Fox et al., 2016), *MASS* (Ripley et al., 2017), and *glmmTMB* (Brooks, et al. 2017). We analysed data on parental care using a zero-inflated binomial model. We used ANOVA models to analyse normally distributed data; that is, female weight change over breeding and mean larval mass at dispersal. We used a quasi-Poisson model to analyse data on female life span and a binomial model to analyse data on larval survival. Note that we did not use a Cox Proportional-Hazards model to analyse female survival as this was not necessary given that we had data on life span of all females, allowing us to compare the life spans of females in the different treatment groups, and because our data did not satisfy the assumption of proportional hazards (Therneau, 2015; χ^2^ = 12.0, P = 0.007). All models included the treatment as a fixed effect with four levels (i.e. infected, immune-challenged, injured and control females). To account for potential effects of brood size on maternal care (Smiseth et al., 2003; Ratz & Smiseth, 2018), we also included brood size at the time of observation as covariate in the model analysing maternal care. We ran pairwise comparisons using a Tukey’s test with the Bonferroni correction whenever the treatment had a significant effect.

To analyse data on gene expression, we first calculated the expression of a gene of interest relative to the reference gene *rpl7* to obtain ΔC_T_ values (Livak & Schmittgen, 2001). We then used ANOVA models to for effects of the experimental treatment on the ΔC_T_ values of each gene. Whenever the treatment had a significant effect on gene expression, we ran pairwise comparisons using a Tukey’s test with the Bonferroni correction.

Among the 245 females, we sacrificed a subset of 59 females to sample hemolymph, of which one was excluded because not enough hemolymph was obtained. Among the remaining females, we excluded 55 additional females from our analysis on maternal care, life span and larval survival because their eggs fail to hatch (N = 10), there were not enough larvae to allocate them a brood (N = 25), the female or the whole brood died before the observation (N = 12), no behavioural data were collected (N = 1), or the heat-kill treatment failed (N = 7). The final sample of the behavioural and life history data included 33 control females, 32 injured females, 33 immune-challenged females, and 33 infected females. Likewise, we excluded 9 broods (control females: N = 4; injured females: N = 3; immune-challenged females: N = 2) from our analysis on mean larval mass at dispersal because no larvae survived to dispersal.

## Results

There was a significant effect of treatment on female life span (figure 1a; χ^2^ = 52.1, df = 3, P < 0.001), which reflected that infected females had an average life span that was 75% shorter than females from any other treatment group (Table 1). There was no significant effect of treatment on the amount of care provided by females (figure 1b; χ^2^ = 6.63, df = 3, P = 0.085), showing that females maintained a similar level of care to control females regardless of whether they were infected, immune-challenged or injured. There was no effect of brood size at the time of observation on maternal care (χ^2^ = 2.62, df = 1, P = 0.105). There was no effect of treatment on mean larval mass at dispersal (Sum Sq = 0.003, df = 3, F = 0.613, P = 0.608) or survival of the larvae until dispersal (χ^2^ = 5.66, df = 3, P = 0.129), suggesting that infected, immune-challenged or injured females maintained a similar level reproductive output to control females. There was no difference in weight change between females in the different treatments (Sum Sq = 174.7, df = 3, F = 1.42, P = 0.239).

**Figure 1.**
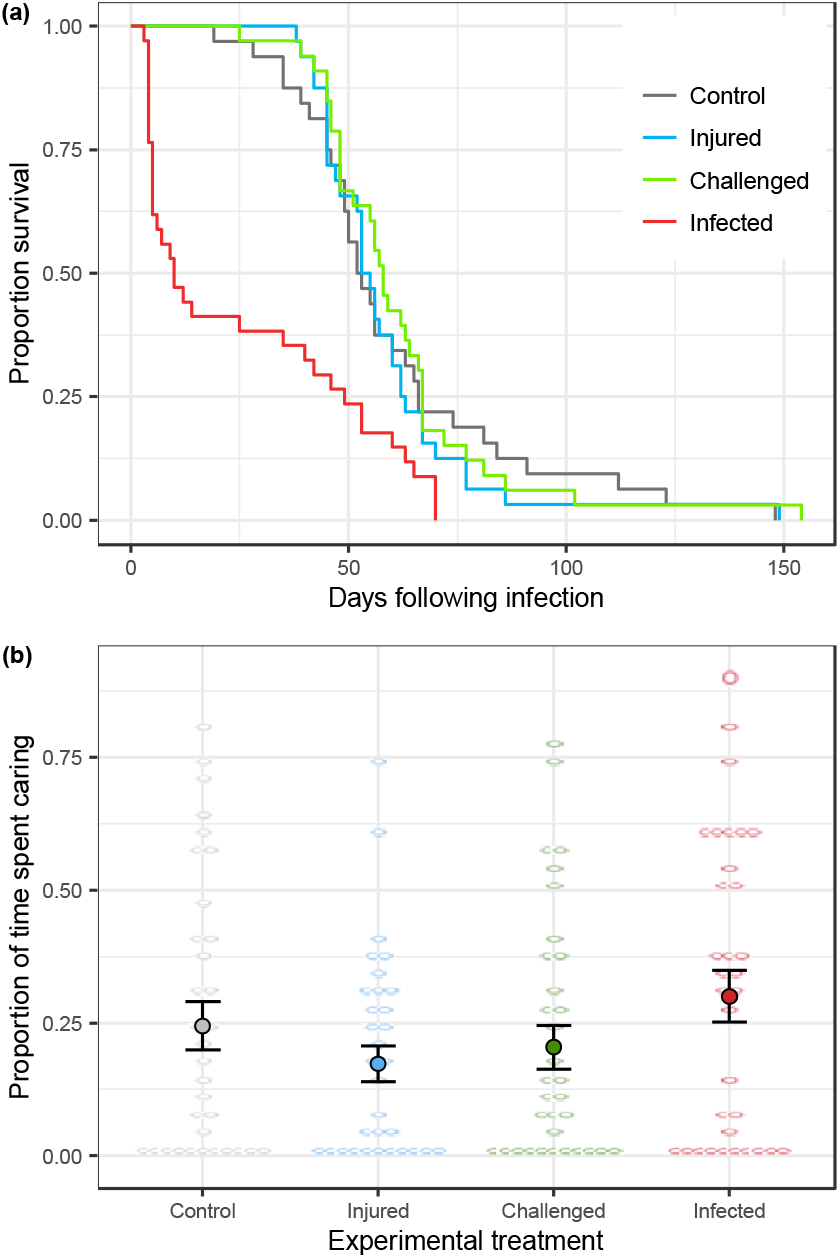
Proportion of females alive over time after the day the treatment was applied (a). Effects of the experimental treatment on maternal care (b). Open circles represent individual data, closed circles and bars represent Means ± SEs.

**Table 1:** Pairwise comparisons between treatments for the post-infection life span. P-values were obtained using Tukey’s HSD test and adjusted using the Bonferroni correction.

| Post-infection life span |  |  |  |  |
| --- | --- | --- | --- | --- |
|  | Estimate | SE | z | P |
| Injured – Control | −0.035 | 0.117 | −0.299 | 0.991 |
| Challenged – Control | 0.014 | 0.115 | 0.123 | 0.999 |
| Infected – Control | −0.866 | 0.148 | −5.83 | <0.001 |
| Injured – Challenged | −0.049 | 0.116 | −0.424 | 0.974 |
| Infected – Injured | −0.831 | 0.149 | −5.57 | <0.001 |
| Infected – Challenged | −0.880 | 0.147 | −5.97 | <0.001 |

Note: Statistically significant P values (<0.05) are shown in boldface.

We next investigated the effects of the experimental treatments on the expression of four immune genes. Treatment had a significant effect on the expression of *coleoptericin-1* (figure 2a; Sum Sq = 780.3, df = 3, F = 42.9, P < 0.0001). The expression of this gene was lower in injured females than in control females (Table 2), lower in immune-challenged females than in injured females (Table 2), and similar in immune-challenged and infected females (Table 2). Treatment also had a significant effect on the expression of *PGRP-SC2* (figure 2b; Sum Sq = 266.7, df = 3, F = 3.47, P = 0.022). The expression of this gene was reduced in injured females compared with infected ones (Table 2), while there was no difference in expression between females in any of the other treatment groups (Table 2). We found no significant effect of treatment on the expression of *attacin-4* (figure 2c; Sum Sq = 45.7, df = 3, F = 1.55, P = 0.211) or *cecropin-1* (figure 2d; Sum Sq = 21.1, df = 3, F = 1.57, P = 0.206).

**Figure 2.**
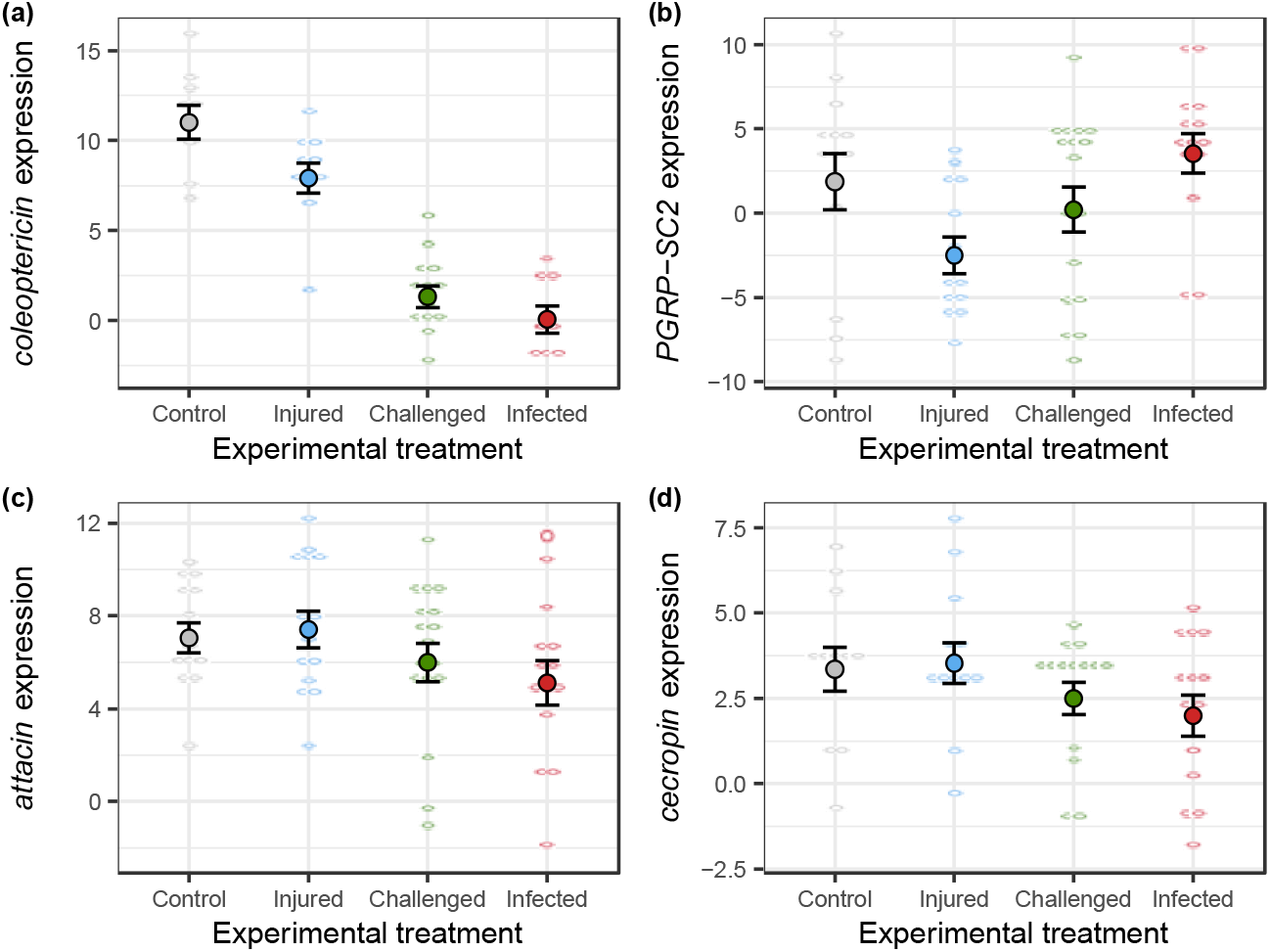
Effects of the experimental treatment on the expression of *attacin-4* (a), *cecropin-1* (b), *coleoptericin-1* (c), and *PGRP-SC2* (d). Open circles represent individual data, closed circles and bars represent Means ± SEs.

**Table 2:**
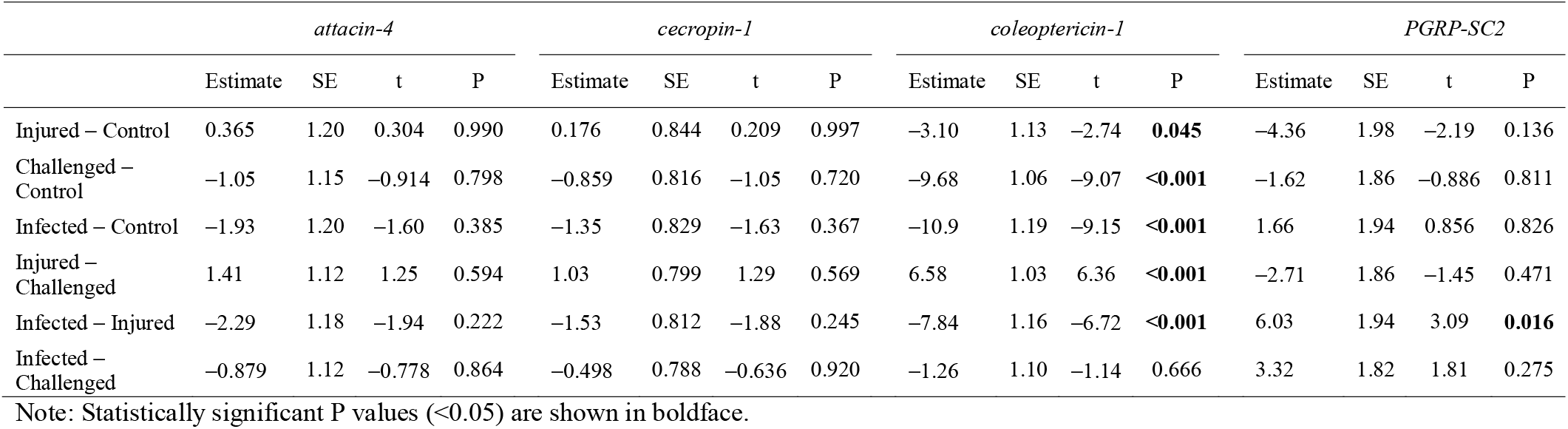
Pairwise comparisons between treatments for the level of gene expression. P-values were obtained using Tukey’s HSD test and adjusted using the Bonferroni correction.

## Discussion

Here we show that infected and immune-challenged females altered their expression of immune genes, and that infected females had a shortened life span compared to other females. Despite the heightened mortality of infected females, we found no evidence for a difference between infected, immune-challenged, injured and control females in their level of care or their reproductive output. Altogether, our findings indicate that infected females maintained their level of care despite indication that they mounted an immune response against the pathogen and clear evidence that the pathogen shortened their life span. This strategy may allow infected females to provide the necessary amount of care to ensure the growth and survival of their offspring but might be detrimental to the parents by increasing their mortality and may potentially even facilitate disease transmission to offspring. Below we discuss the broader implications of these findings to our understanding of the effects of infection on parental behaviour and social interactions between caring parents and their dependent offspring.

As expected, we found that infected females altered their expression of immune genes and had a considerably shortened life span, confirming that infection with *Serratia marcescens* had the intended effect of triggering an immune response and making infected females sick. Immune-challenged females showed a similar change in the expression of immune genes as infected females, but suffered no corresponding reduction in their life span. Thus, our results confirm that the shortened life span of infected females was caused by the pathogen rather than being a by-product of females mounting an immune response. Taken together, our results confirm that *Serratia marcescens* is a potent pathogen in *N. vespilloides*. We are not aware of any prior studies on *N. vespilloides* reporting elevated mortality as a result of an infection, which may reflect the difficulty in establishing experimental infections in this species. This may reflect that this species breeds on decomposing carcasses, which means they regularly will be in close contact with potential pathogens (Jacobs et al., 2014; Wang & Rozen, 2018). Our study species might thus be resistant to a wide variety of bacterial strains, such as *Bacillus subtilis* (Reavey et al., 2015), *Pectobacterium carotovorum*, *Pseudomonas aeruginosa*, *P. entomophila*, or *S. marcescens* at low doses and concentrations (Ratz et al., unpublished data) that are pathogenic in many others insect species. Our results show that, as long as *S. marcescens* is injected in relatively high dose and concentration, it can successfully establish an infection in *N. vespilloides,* activate the immune system, and greatly increase mortality.

Our main finding was that infected females maintained their level of care and their reproductive output, despite that these females had mounted an immune response and were suffering negative fitness consequences of infection as indicated by their shortened life span. Our results suggest that infected females maintained their level of care at the expense of allocating more resources towards immunity. Our results are similar to those of a recent study on the amphipods *Crangonyx pseudogracilis* and *Gammarus duebeni* (Arundell et al., 2014). In this study, infection by a microsporidian did not affect brood care behaviour or the duration of brooding of females. By maintaining their level of care, infected females may ensure that offspring receive the necessary amount of care and produce offspring with a similar survival and body size as offspring of uninfected females. This strategy might allow infected females to maintain their reproductive output (Arundell et al., 2014), but might come at a cost in terms of reduced survival and future reproductive success. Burying beetles can produce multiple broods (Creighton et al., 2009) and tend to gain mass during first reproductive, which is positively correlated with life span (Gray et al., 2018). Our results suggest that infected females would have lower fitness because it seems unlikely that the infected females in our study could reproduce again. The reason for this is that approximately 60% of infected females had died by 17 days after the infection (compared with 0% of control females; figure 1a). Thus, many infected females had died before they would have been able to produce an additional brood. In order to breed again, females must first remain with the current brood until larvae complete their development, which would take about 7 days (Smiseth et al., 2003; 2005). They then need to search for and secure a new carcass, which are thought to be rare (Scott, 1998), and produce eggs and care for the new brood, which would take another 10 days (Ford & Smiseth, 2017). An alternative explanation for our results is that infected females perceived their chance to survive and reproduce again to be very low, and that they therefore maintained a high level of care as a terminal investment response (Williams, 1966). This is suggested by other studies in the species reporting high reproductive output in response to immune-challenges (e.g. Cotter et al; 2010; Reavey et al., 2014; Reavey et al., 2015; Farchmin et al., 2020). We found no evidence for an increase in reproductive investment as would be expected under terminal investment. However, this may reflect that infected females were simply not able to increase their level of care. We would have expected immune-challenged females, exposed to pathogen cues but not infected, to be able to increase care given that they did not show any evidence of shortened life span. We did not find such a response in immune-challenge females. Thus, we suggest that, rather than mounting a terminal investment response, infected females maintained their level of care to provide the necessary amount of care to ensure offspring growth and survival, which might come at a cost to females in terms of reduced survival.

Our finding that infected females maintained their level of care also shows that infections do not necessarily induce sickness behaviour. Infections are often associated with a reduction in the host’s social interactions (Hart, 1988; Kelley et al., 2003), which was not the case in our study as there was no evidence for a reduction in maternal care. Infected hosts often show reduced social interactions (Vale et al., 2018), which may be the result of lethargy (i.e., reduced activity levels) of the host associated with sickness (Adelman & Martin, 2009), the host actively avoiding costly social interactions (Sah et al., 2018; Lopes et al., 2016), uninfected individuals avoiding an infected host (Curtis, 2014), or the pathogen manipulating the host’s behaviour (Moore, 2002; Hughes et al., 2012). Yet this reduction in social behaviour is not always observed, depending on the social context (Lopes et al., 2012; Adamo et al., 2015), and parents that are sick might maintain their level of care and interactions with offspring (Stockmaier et al., 2020). Because parental care and parent-offspring interactions can have a large impact on the reproductive output of organisms, we propose that infected parents might prioritise their allocation in reproduction by maintaining necessary care and social interactions with their offspring. In species with biparental care, infected females might be able to reduce their level of care (and thereby increase their immune response) without harming their offspring if the male parent compensate for the reduction in female care. If so, male compensation could temper the negative effect of infection on female life span. Thus, we encourage future studies to compare the responses of infected females in the contexts of biparental care and uniparental care.

Our last finding was that females from our different treatment groups showed different levels of expression in two immune genes (i.e. *coleoptericin-1* and *PGRP-SC2*), while there was no difference in the expression of two other immune genes (i.e. *attacin-4* and *cecropin-1*). The expression of *coleoptericin-1,* a gene involved in personal immunity (Jacobs et al., 2016; Parker et al., 2015), was lower in immune-challenged and infected females than in injured and control females. In contrast, the expression of *PGRP-SC2,* a gene involved in social immunity (Parker et al., 2015; Ziadie et al., 2019), was higher in infected females than in injured females. Given that there was no difference in immune gene expression between immune-challenged and infected females, it seems unlikely that the pathogen supressed the immune system in our study species. Instead, these results might reflect immune responses to the presence of a pathogen or, in the case of immune-challenged females, to the presence of cues from a potential pathogen. Thus, our finding that infected females had lower personal immunity and higher social immunity points towards a shift in investment towards current reproduction. This suggests that infected and immune-challenged females maintained their investment in social immunity that benefit larval survival, which would be in line with the idea that infected females overall sought to maintain their allocation towards current reproduction. However, we urge caution in interpreting our results given that our study focused on the expression of four immune genes, which might not reflect the immune response as a whole.

Our findings have important implications for our understanding of parental behaviour under the risk of infection by showing that infected females maintained a high level of care despite that infections could expose their offspring to the pathogen. Thus, our results show that the level of care is remarkably stable in response to infection, despite evidence that parents often show a great amount of plasticity in response to other environmental factors, such as resource abundance and the presence of competitors and infanticidal conspecifics (Smiseth & Moore, 2002; Hopwood et al. 2015; Georgiou Shippi et al. 2018). Furthermore, behavioural plasticity represents the first mechanism of immunity (Schaller, 2006; Schaller & Park, 2011; Kiesecker et al., 1999) and might allow infected individuals to reduce the risk of transmission to close kin, such as offspring (Shakhar & Shakhar, 2015; Shakhar, 2019). Our study found no evidence that females transmitted the pathogen to their offspring given that we found no evidence that larvae of infected females had lower survival than larvae of other females. Nevertheless, we urge future studies to consider the potential consequences of disease transmission by caring parents to their offspring (Chakarov et al. 2015). For example, infected parents might be expected to maintain their level of care in situations where the risk of females passing on the pathogen to their offspring is low. In contrast, infected parents might reduce their level of care in situations where the risk of females passing on the pathogen to their offspring is high and where the offspring are not completely dependent on their parents.

In summary, our study shows that infected females maintained their level parental care and reproductive output despite mounting an immune response and suffering from greater mortality. Our results demonstrate that parental care, which is generally highly flexible, can remain robust and stable in response to pathogenic infections. Our results suggest that infected females maintain their current reproductive success over survival, which could ensure that offspring receive the necessary amount of care. Our findings stress the need for more studies on infection in species where parents care for and interact with their offspring, as parental care is a fundamental social interaction in all birds and mammals as well as some amphibians, fishes and arthropods and as it can have contradicting effects by buffering against environmental hazards on the one hand and providing a potential route for disease transmission on the other hand.

## Acknowledgments

We thank the City of Edinburgh Natural Heritage Service for permission to collect beetles in their reserve at the Hermitage of Braid and Blackford Hill Local Nature Reserve. We also thank Jon Richardson for assistance with maintaining the laboratory population, Arun Prakash, Sarah Reece, Saudamini Venkatesan, Ferghal Waldron and Michelle Ziadie for suggestions and useful advice on the experimental set up, and Eevi Savola for helpful discussions regarding the analysis of gene expression data. T.R. was supported by the Darwin Trust of Edinburgh.

## Notes

### Competing Interest Statement

The authors have declared no competing interest.

